# DDR1 regulates RUNX1-CBFβ to control breast stem cell differentiation

**DOI:** 10.1101/2024.02.21.581255

**Authors:** Colin Trepicchio, Gat Rauner, Nicole Traugh, Meadow Parrish, Daniel E.C. Fein, Youssof Mal, Charlotte Kuperwasser

**Author notes:** **Correspondence should be addressed to:** Charlotte Kuperwasser.

## Abstract

The human breast is complex and comprised of multi-lineage and multi-structural elements. Recent work has shown that epithelial stem and progenitor cells use the collagen receptor Discoidin Domain Receptor 1 (DDR1) for differentiation into both basal and luminal cell lineages, which together are necessary for both ductal and alveolar morphogenesis. We developed a next-generation single cell derived organoid model that generates miniaturized breast tissue, to study how single stem cells can give rise to multiple cell types and compound tissue structures. We show that DDR1 activation triggers stem cell differentiation via RUNX1 in turn driving multilineage differentiation as well as complex ductal-lobular development. Mechanistically, DDR1 affects the interaction and expression of RUNX1 and its cofactor CBFβ thereby regulating its activity. Together, these findings contribute to the current understanding of how the extracellular matrix component within the stem cell niche drives organogenesis and tissue regeneration.

## Introduction

Epithelial organogenesis, morphogenesis, and differentiation are core processes of developmental biology, underlying the specification, expansion, and differentiation of stem cells into organized multicellular tissues (Thomson, 1917; Waddington, 1941, 1957). In the case of the human breast, which is characterized by both its complex ductal and lobular anatomy and its multilineage epithelium, the process by which these structures arise from stem cells and how they relate to each other during development has been difficult to study. In addition, how stem cell differentiation and tissue morphogenesis is controlled in a highly regenerative tissue such as the breast is important to understand as defects in these processes underlie the formation of breast cancer (Arendt et al., 2010, 2014; Ferraro et al., 2010; Hutson et al., 1985; Petersen and Polyak, 2010; Russo and Russo, 2004; Wellings et al., 1975; Woodward et al., 2005).

The human breast is a dynamic tissue that undergoes several developmental phases, starting during embryogenesis with organogenesis, then continuing postnatally at puberty, again during pregnancy, lactation, and finally regressing during post-lactation involution. The breast actively regenerates during each menstrual and pregnancy cycle until ultimately atrophying and losing its regenerative capacity at menopause (Arendt et al., 2014; Rauner et al., 2023; Russo and Russo, 2004; Williams and Daniel, 1983). Unlike rodents, human breast tissue is composed of 15 to 20 lactiferous ducts which are responsible for transporting milk from the lobules to the nipple. Lobules with differing levels of complexity are composed of many terminal ductal lobular units (TDLUs) that consist of terminal ducts culminating in grape-like clusters of alveoli (Arendt et al., 2014; Hutson et al., 1985; Russo and Russo, 2004; Wellings et al., 1975; Williams and Daniel, 1983). A single lobule can have anywhere from 10 to 100 TDLUs, each composed of both luminal epithelial cells that line the inside of the ducts and alveoli and basal/myoepithelial cells that lie outside the luminal cells and are in direct contact with the basement membrane (Arendt et al., 2014; Hutson et al., 1985; Russo and Russo, 2004; Wellings et al., 1975; Williams and Daniel, 1983; Woodward et al., 2005). During development, both luminal and myoepithelial cells are derived from bipotent stem cells that give rise to unipotent luminal and basal progenitors, which in turn serve as precursor reservoirs for mature luminal and myoepithelial cells in the adult tissue, respectively (Arendt et al., 2014; Hutson et al., 1985; Wellings et al., 1975; Williams and Daniel, 1983; Woodward et al., 2005)

The ability of breast epithelial stem cells to create complex structural components, such as ducts and alveoli is governed by a combination of intrinsic cell programming and cell-cell interactions, as well as extrinsic signals from the microenvironment (Williams and Daniel, 1983; Woodward et al., 2005). Mouse models have been traditionally used to study mammary stem cells, development, and morphogenesis (Bresslau, 1920; Holen et al., 2017), however, there are inherent differences between species that limit the direct translation of findings to humans (Bresslau, 1920; Gregory, 1910; Holen et al., 2017; Russo and Russo, 2004). Traditional models including 2D cell cultures and 3D organoid and mammosphere models have also provided valuable insights in studying breast stem cells and epithelial differentiation (Azimian Zavareh et al., 2022; Kim et al., 2004). However they fall short in capturing the complexities of human breast tissue organogenesis and morphogenesis (Kim et al., 2004).

Recently, we reported the creation of a next-generation 3D organotypic model with advanced representation of complex glandular human TDLU architecture and function (Rauner et al., 2021; Sokol et al., 2016). We found that human breast TDLU organoids form in response to collagen signaling through the activation of discoidin domain receptor 1 (DDR1) (Rauner et al., 2021). DDR1 activation by collagen gives rise to basal progenitors and luminal cells, which drive alveolar budding, branching, and the formation of complex TDLU organoids^16^. In this study, we sought to study how DDR1 signaling in single cells leads to the creation of these complex multi-structural tissues. In doing so we identified RUNX1 as a necessary transcriptional regulator linking DDR1 signaling with bi-potent stem cell differentiation that in turn is required for tissue morphogenesis. These findings along with mutational data suggest that disruption of this signaling axis could have important implications in the biology of breast cancer.

## Results

### Live imaging of single cells reveals the complex and dynamic nature of TDLU organoid formation and morphogenesis

To study stem cell dynamics and tissue morphogenesis, we seeded single human breast epithelial cells into 3D hydrogels. This results in the formation of complex, multilayered and heterogeneous breast TDLU organoids through a series of well-coordinated stages (Fig. 1A) (Rauner et al., 2021; Sokol et al., 2015, 2016). We combined this method with high-speed point scanning confocal microscopy, to visualize organogenesis from a single bipotent stem cell in real time (Fig. 1B, Supplemental Video 1). We observed that TDLU organoid formation takes ∼18-21 days to complete and proceeds through 4 distinct stages: Induction, Patterning, Morphogenesis & Maturation. Within the first 4 days after seeding, stationary single stem cells are induced to proliferate and exit the bipotent stage (Fig.1B, Supplemental Video 1) (Rauner et al., 2023). Shortly thereafter, between days 5-8, stem cell progeny exhibit dynamic cell movements where they travel along an emerging branch prior to the formation of a cohesive tissue (Fig.1B, Supplemental Video 1). Epithelial cells continue to divide and invade the matrix, in a manner reminiscent of tissue patterning, clearing room for the organoid to develop primary ducts. Around day 9 basal progenitor cells begin the process of morphogenesis whereby ducts elongate and alveolar buds appear (Fig.1B, Supplemental Video 1). This leads to the formation, maturation and differentiation of organoids that are anatomically equivalent to human breast TDLUs (Fig. 1C i-v). These structures are histologically normal with an inner layer containing mature E-cadherin expressing luminal cells (Fig. 1C i, iii, v) and external CK14^+^ myoepithelial cells (Fig. 1C i, iv, v). Since TDLU organoids are derived from a single cell (Supplemental Video 1) (Rauner et al., 2023), and the resulting complex tissue is comprised of both ducts and alveoli as well as both basal and luminal cell lineages, (Fig. 1B,C), this parent–progeny relationship between the precursor cell and the TDLU demonstrates they are facultative multilineage stem cells (Ferraro et al., 2010; Petersen and Polyak, 2010; Rauner et al., 2023; Woodward et al., 2005).

**Figure 1:**
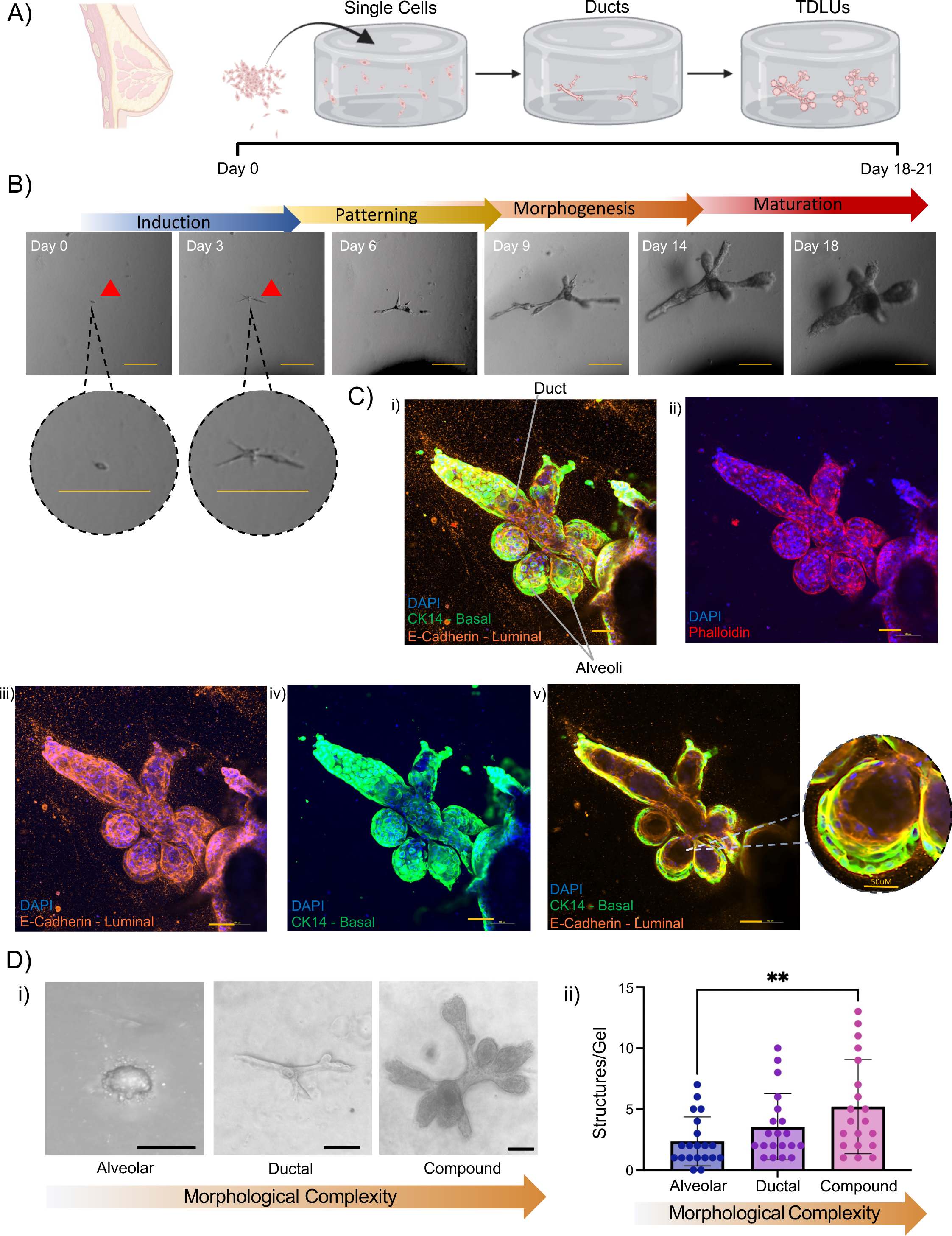
Phenotypic characterization of patient-derived single cell hydrogel methodology. A) Schematic representation and summary of human breast organogenesis in a 3D hydrogel TDLU organoid model. B) Timeline with brightfield images of TDLU formation from single cells, starting on day 0 immediately after hydrogel polymerization to the formation of complex ductal-lobular structures from day 18 onward. Four phases of organogenesis showing induction, patterning, morphogenesis, and maturation. Scale bar = 100 μm. C) i-iv) Immunostaining of a Three-dimensional Maximum Intensity Projection (MIP) TDLU organoids, at day 21 of development, stained with phalloidin (red) for actin cytoskeleton, with CK14 (green), E-cadherin (orange) for cell lineage, and with DAPI (blue) for nuclei. Scale bars = 100 μm v) Two-dimensional cross-section of MIP (i) with offset highlighting a lobule showing distinct CK14^+^ (green) and E-Cad^+^ (orange) layers with an interacting layer displaying yellow. Scale bar = 100 μm, scale bar of offset = 50 μm. D) Representative brightfield images (i), and quantification (ii) of 3D organoid morphologies from 5 primary patients. Mean ± SD (n= 4 gels/primary patient samples). Scale bar = 100 μm.

Notably, not all organoids originating from single cells form highly compound TDLUs. Rather, some remain in a simpler state, categorized as alveoli only or duct only. Alveolar organoids are characterized by the formation of rudimentary acini or clusters of cells that lack a defined architecture, while duct only organoids are characterized by the formation of elongated and branched structures but lack alveoli. In contrast, complex TDLU organoids are compounded structures containing both ducts and varying numbers of alveoli at their terminal regions (Fig. 1D). A quantitative assessment of these organoid types across five patient samples revealed that they skew towards the formation of these compound TDLU structures, averaging 11 total organoids per 1000 seeded cells, with about of 2.35 Alveolar, 3.35 Ductal, and 5.2 compound TDLU structures in each gel (Fig. 1D).

### Inhibition of DDR1 restricts stem cells to a bipotent state and blocks alveologenesis

Using this approach, we examined the effects of DDR1 during development. DDR1, a receptor tyrosine kinase activated by collagen binding, has recently been shown to be required for breast organoid development (Rauner et al., 2021). Single epithelial breast cells were treated with a small molecule inhibitor of DDR1 (DDR1i) on day 0 after seeding to study its function in single stem cells (Fig. 2A). A total lack of TDLU formation was observed when single stem cells were treated with DDR1i during induction as well as a significant reduction in the total number of organoids that formed (Fig. 2B, Supplemental Fig. 1A). Moreover, any organoids that managed to form in the presence of DDR1i at induction displayed noticeable abnormalities. Staining of these disorganized, rudimentary structures with E-cadherin and CK14 revealed that the cells within the structures were double positive for both luminal and basal markers suggesting that were trapped in a bi-potent state (Fig. 2C).

**Figure 2:**
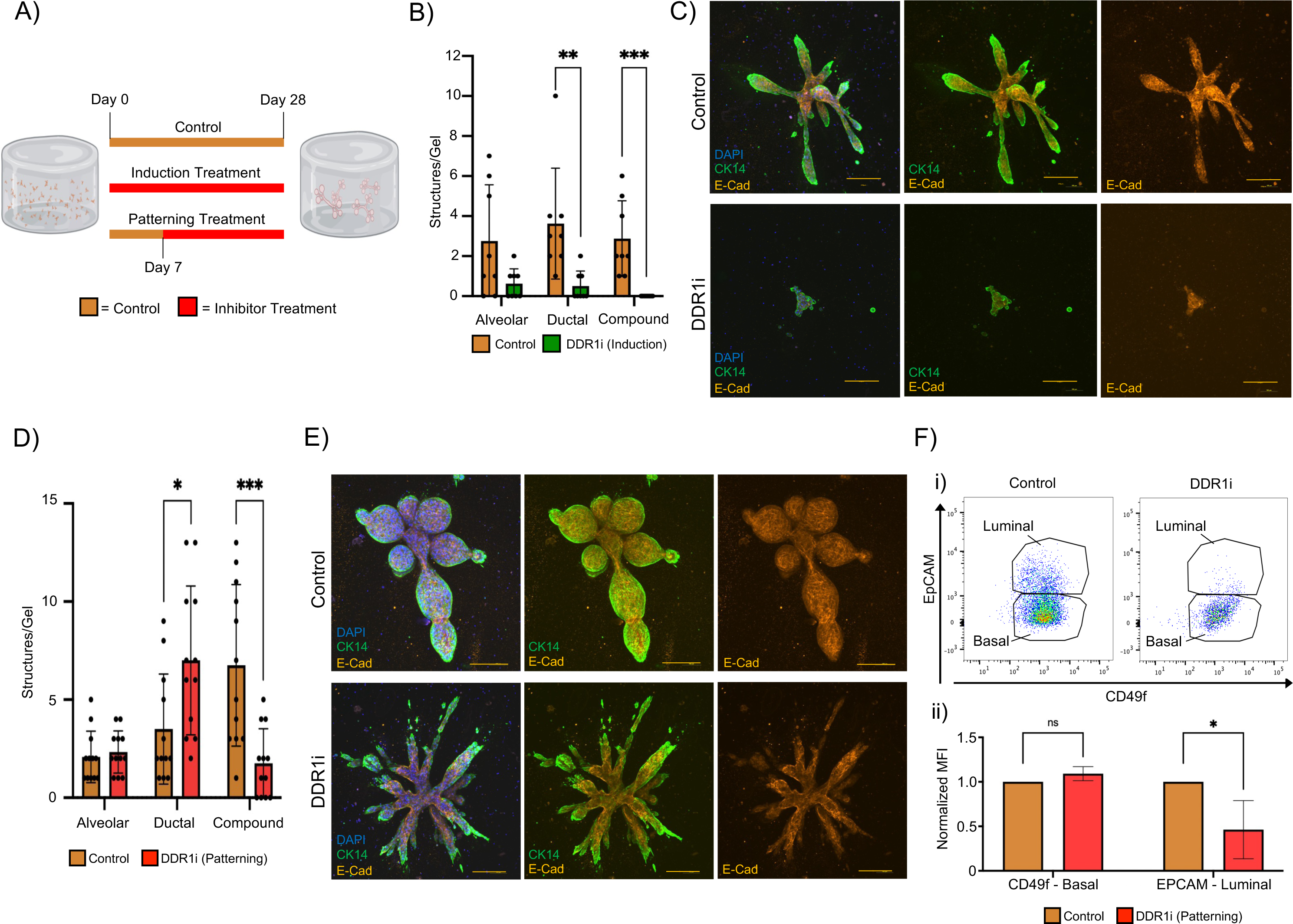
Effects of DDR1 inhibition during induction and patterning from single cell derived miniature TDLU organoids. A) Schematic of the strategy to test the effects of DDR1i on breast TDLU organogenesis. Inhibition during induction began with DDR1i treatment starting at day 0 and concluded when control structures were fully formed no later than day 28. Inhibition during patterning started with DDR1i beginning day 7 and concluding around day 28. B) Quantification of the types of organoids that formed following DDR1i treatment during induction. Data presented as Mean ± SD (n= 4 gels/primary patient samples). C) Representative immunofluorescence staining of organoids from control or DDR1 inhibitor-treated gels, with treatment initiated during the induction phase of organoid formation. CK14 (green), and E-Cadherin (orange), and DAPI (blue) staining. Scale bar = 200 μm. D) Quantification of the types of organoids formed in DDR1i treated cultures beginning during patterning. Data presented as Mean ± SD (n= 4 gels/primary patient samples). E) Representative immunofluorescence staining of organoids from control or DDR1i treated gels initiated during the patterning phase of organoid formation. CK14 (green), and E-Cadherin (orange), DAPI (blue) staining. Scale bar = 200 μm. F) Representative flow cytometry plot analysis of basal (CD49f) and luminal (Epcam) cells (FACS) (i) and quantification of mean fluorescent intensity (ii) from primary patient samples cultured in 3D, treated with DDR1i during patterning (n=3 primary samples, and values are expressed as Mean ± SD). Statistical significance was determined via multiple t-tests, with significance levels indicated as follows: *p-value < 0.05, **p-value < 0.01, ***p-value < 0.001, ****p-value < 0.0001.

We next examined how DDR1 functions later in development after stem cell induction. 3D cultures were next treated with DDR1i beginning on day 7 (Fig. 2A), to study the role of DDR1 during patterning. While the total number of organoids that formed was similar to the control, there was also a failure to form complex ductal-alveolar TDLUs in the presence of DDR1i during patterning (Fig. 2D, Supplemental Fig. 1B). Rather, organoids that formed in the presence of DDR1i remained simple and were primarily restricted to ductal-only structures.

Staining of these organoids with E-cadherin and CK14 revealed lineage specification with the development of both ductal luminal and basal cells (Fig. 2E). This suggested that inhibition of DDR1 during patterning does not block stem cell differentiation but rather affects tissue patterning and formation of alveoli necessary to form TDLUs.

The capacity to undergo alveolar morphogenesis requires coordinated expansion and differentiation of luminal cells (Rauner et al., 2021; Rodilla et al., 2015; Yamaji et al., 2009). Thus, we examined whether the lack of alveolar morphogenesis in response to DDR1i, might be due to a failure of luminal cell expansion. Luminal (EpCAM^high^) and basal (EpCAM^neg/low^/CD49f^pos^) cells were assessed by flow cytometry in organoids that formed in the presence or absence of DDR1i during patterning. A significant reduction in the number of luminal cells was found in DDR1i treated organoids (Fig. 2F), consistent with the notion that the failure to form alveoli is due to a failure of luminal cell expansion (Rauner et al., 2021; Rodilla et al., 2015; Yamaji et al., 2009). Together, these findings along with prior studies (Rauner et al., 2021) demonstrate that DDR1 plays a crucial role in driving the specification and differentiation of stem cells during induction but also the expansion of luminal cells during patterning.

### DDR1 regulates RUNX1 expression and transcriptional activity

Studies have suggested that RUNX1 is essential for breast stem cells to exit the bipotent state and commit to a specific lineage (Hong et al., 2017; Rose et al., 2020; Sokol et al., 2015). Given that DDR1i also blocked stem cell lineage specification and differentiation, we examined scRNA-seq for RUNX1 expression in organoid cultures treated with DDR1i. Organoids were grown for 14 days and treated continuously with DDR1i or treated with DDR1i for 12 days followed by release from inhibition for two additional days (DDR1r) (Fig. 3A) (Rauner et al., 2021). In the absence of DDR1i, RUNX1 is heterogeneously expressed in various cell types including bipotent stem cells, basal cells, and mature luminal cells (Fig. 3B). The highest levels of RUNX1 are expressed in bipotent progenitor cells and basal cell types, with lower levels expressed in luminal cells. Interestingly, DDR1 is also expressed in similar cell populations including bipotent progenitor and basal cells with lower levels in luminal cells (Supplemental Fig. 2A).

**Figure 3:**
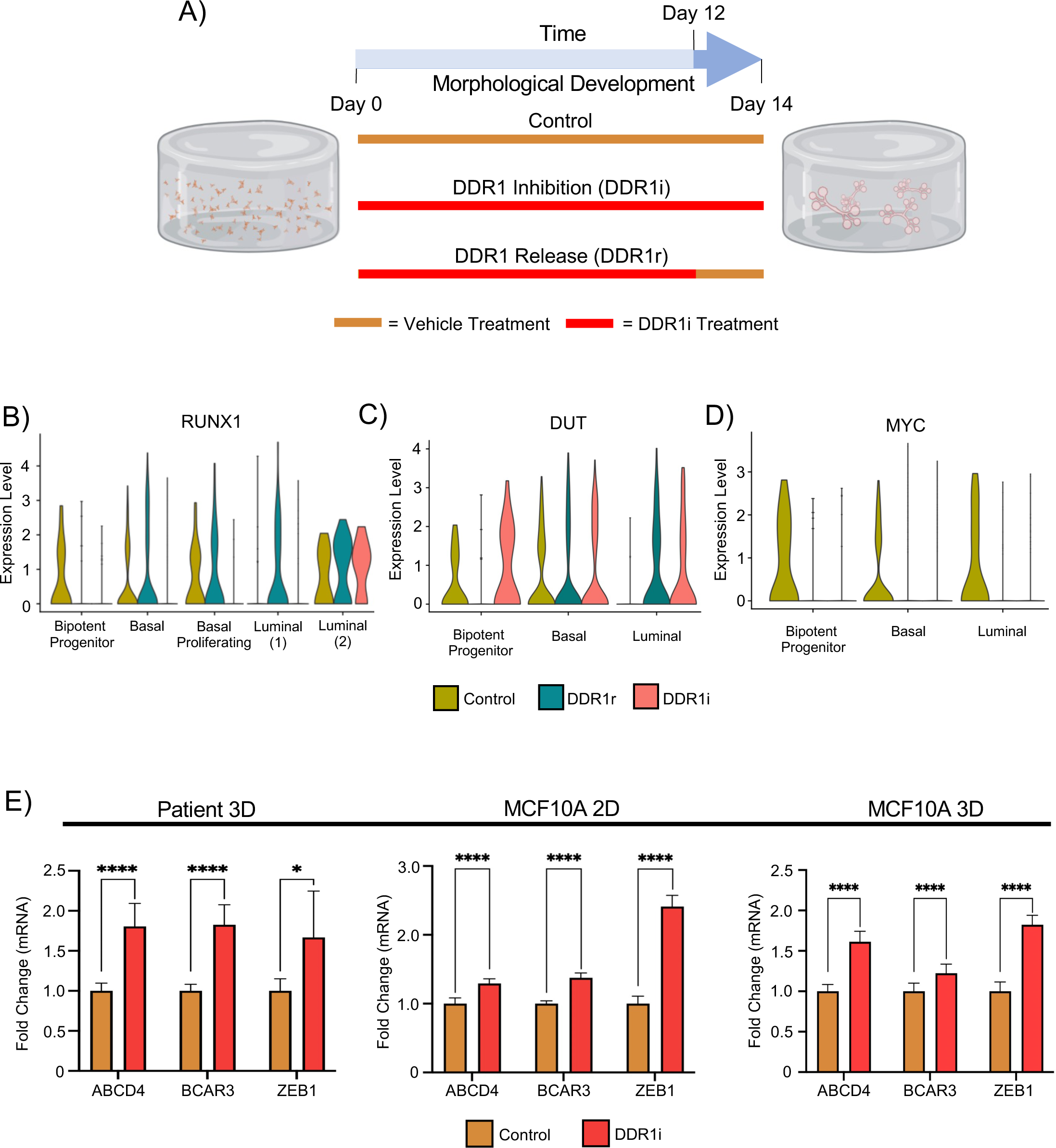
Effects of DDR1 inhibition on RUNX1 and RUNX1 target genes. A) Schematic representation of DDR1 inhibitor time course design for scRNA-seq. DDR1i treatment starting on day 0 and concluding on day 14, and DDR1r treatment with DDR1i initiation on day 0 and cessation on day 12, with the last two days free from inhibition. B-D) Violin plots showing the distribution of RUNX1, DUT, and MYC expression in primary tissue organoids under control, DDR1r or DDR1i conditions. E) Compilation of qRT-PCR quantification data derived from three primary patient samples cultured in a 3D environment, MCF10A cells grown in 2D or MCF10A cells grown in 3D, comparing RUNX1 target gene response to DDR1 inhibition. Values expressed as Mean ± SD. Statistical significance was determined via multiple t-tests, with significance levels indicated as follows: *p-value < 0.05, **p-value < 0.01, ***p-value < 0.001, ****p-value < 0.0001.

Notably, DDR1i led to a substantial downregulation of RUNX1 expression in both basal cell populations and bipotent progenitor cells. While this downregulation was reversible in basal cells, as the expression of RUNX1 was restored once DDR1i was removed (DDR1r) (Fig. 3B), bipotent progenitors showed an irreversible downregulation of RUNX1, suggesting that bipotent progenitors and basal cells likely have different regulatory mechanisms governed by DDR1. Interestingly, Luminal 2 cells, the only cell type in the breast in which DDR1 is not expressed is also the only cell type where DDR1i does not affect RUNX1 expression (Fig. 3B, Supplemental Fig. 2A).

Given that RUNX1 expression levels were sensitive to DDR1 activity, we also examined whether RUNX1 target gene expression might be responsive to DDR1i. Indeed, DDR1i caused differential expression in several direct RUNX1 target genes including DUT, an essential nucleotide metabolism enzyme seen upregulated across breast cancers (Davison et al., 2021), and MYC (Fig. 3D, E, Supplemental Fig. 2B-F). The responsiveness of RUNX1 to DDR1i was specific as expression of other members of the RUNX family (RUNX2, RUNX3) (Supplemental Fig. 2G, H) or its essential cofactor Core binding factor beta (CBFβ) (Supplemental Fig. 2I) were not affected by DDR1i.

To further validate these findings, we treated three additional primary organoid cultures and assessed mRNA expression of additional RUNX1 target genes to DDR1i. While patient heterogeneity in response to DDR1i was observed (Supplemental Fig. 2J, panels i-iii), RUNX1 target genes were similarly affected by DDR1i (Fig. 3E). We also examined the expression of these RUNX1 target genes in MCF10A and MCF10F cells, normal immortalized breast cell lines that maintain populations of basal and luminal lineages (Tait et al., 1990). Cultured in 2D and stimulated with collagen and DDR1i, MCF10A (Fig. 3J) and MCF10F (Supplemental Fig. 2K) cells showed a significant change in mRNA expression of these RUNX1 target genes. MCF10A cells grown in 3D also showed a significant change in RUNX1 target gene expression (Fig. 3K). Taken together, this data shows that DDR1 activity can modulate RUNX1 expression and its target genes.

### RUNX inhibition phenocopies DDR1 inhibition

Given that DDR1 regulates the expression of RUNX1 and its target genes, and that RUNX1 is necessary for stem cell differentiation (Sokol et al., 2015), we next examined whether inhibition of RUNX1 might also affect organoid formation in a manner akin to DDR1i. To this end, growing organoid cultures were treated with a pan-RUNX inhibitor AI-10-104 (RUNXi), that blocks RUNX activity by preventing binding to CBFβ (Illendula et al., 2016), on day 0, prior to stem cell exit from bipotency, or on day 7 during progeny patterning. Similar to DDR1i treatment, when single stem cells were treated with RUNXi during induction, we observed structural arrest in alveolar and ductal stages, leading to a complete loss of mature TDLU organoid formation, similar to DDR1i, but not a reduction in the total number of organoids that formed (Fig. 4A Supplemental Fig. 3A). The colonies that did develop in the presence of RUNXi were immature, disorganized, and comprised of cells double positive for both luminal and basal markers, consistent with stem cells being trapped in a bi-potent state (Fig. 4B) (Sokol et al., 2015).

**Figure 4:**
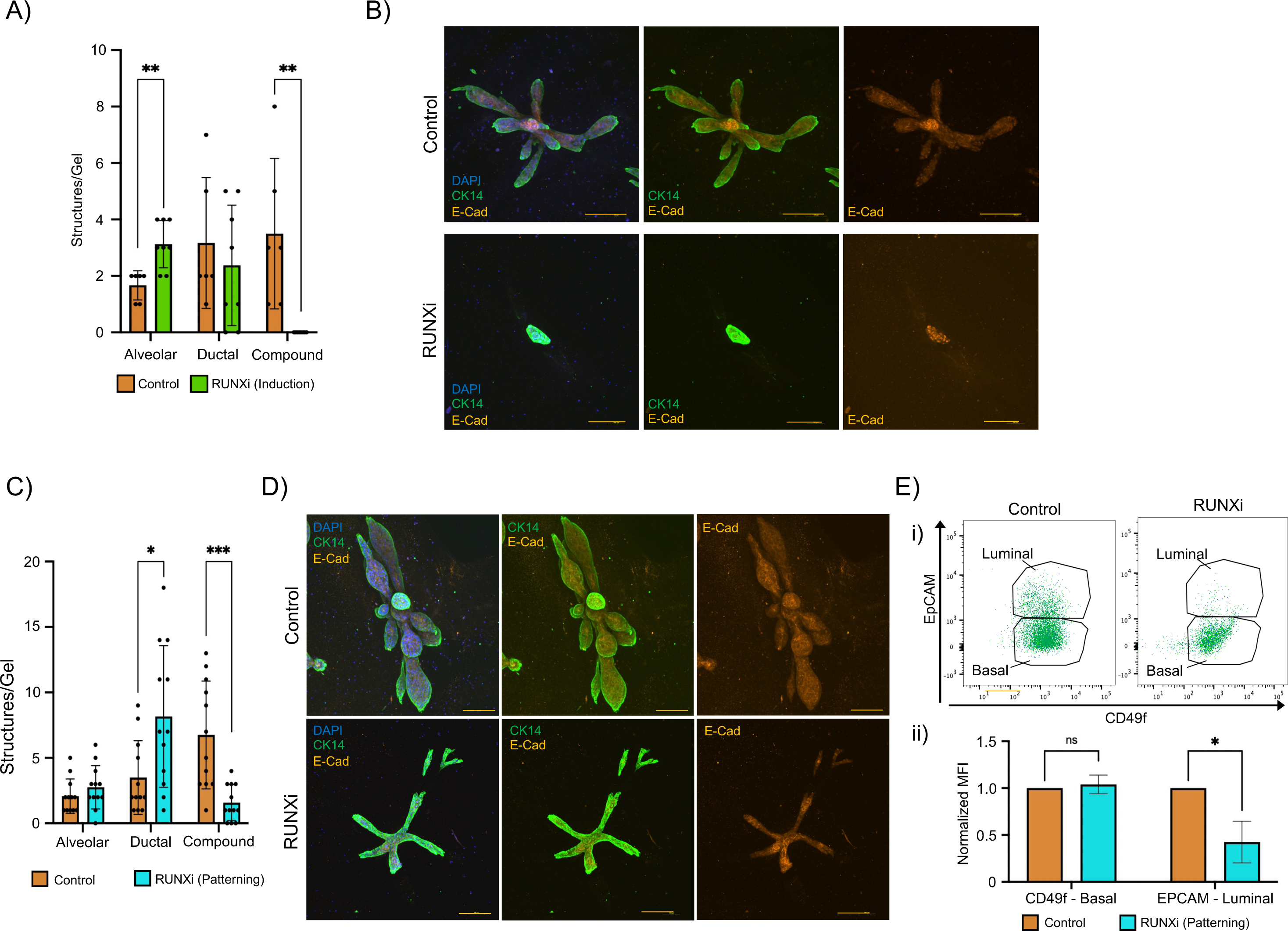
Effect of RUNX inhibition during induction and patterning from single cell derived miniature TDLU organoids. A) Quantification of the types of organoids that formed following RUNXi treatment during induction. Data presented as Mean ± SD (n= 4 gels/primary patient samples). B) Representative immunofluorescence staining of organoids from control or RUNX inhibitor-treated gels, with treatment initiated during the induction phase of organoid formation. CK14 (green), and E-Cadherin (orange), and DAPI (blue) staining. Scale bar = 200 μm. C) Quantification of the types of organoids that formed following RUNXi treatment during patterning (n= 4 gels/primary patient samples). Data presented as Mean ± SD. D) Representative immunofluorescence staining of organoids from control or RUNX inhibitor-treated gels, with treatment initiated during the induction phase of organoid formation. CK14 (green), and E-Cadherin (orange), and DAPI (blue) staining. Scale bar = 200 μm. E) Representative flow cytometry plot analysis of basal (CD49f) and luminal (Epcam) cells (FACS) (i) and quantification of mean fluorescent intensity (ii) from primary patient samples cultured in 3D, treated with RUNX inhibitor during patterning (n=3 primary samples). values are expressed as Mean ± SD. Statistical significance was assessed using multiple t-tests, with significance levels indicated as follows: *p-value < 0.05, **p-value < 0.01, ***p-value < 0.001, ****p-value < 0.0001.

Like DDR1i during patterning, RUNXi treatment during patterning resulted in the loss of complex TDLU organoid formation (Supplemental Fig. 3B). Instead, organoids that formed were composed primarily of simple ducts, although the total number of organoids was not affected by RUNXi (Fig. 4D, Supplemental Fig. 3C) pointing towards a blockage in tissue patterning. Staining of the RUNXi organoids with E-cadherin and CK14 also showed that, like DDR1i, lineage specification with the development of both ductal luminal and basal cells was present despite the failure to form alveoli (Fig. 4E). Together, these results imply that inhibiting RUNX during epithelial patterning does not block the differentiation of bipotent stem cells but rather abrogates the capacity of luminal and basal cells to organize into complex structures, mimicking structures generated with DDR1i at this stage.

We also examined whether the lack of alveolar morphogenesis in RUNXi treated organoids, might be due to differences in cell types present within the structures. Notably, a significant reduction in the number of EpCAM^high^ luminal cells was found in RUNXi organoids (Fig. 4F) consistent with the notion that the failure to organize alveoli is due to a failure of luminal cell expansion (Rauner et al., 2021; Rodilla et al., 2015; Yamaji et al., 2009). Collectively, these results show that RUNX also plays a crucial role in driving the differentiation of bi-potent stem cells, as well as in the expansion of luminal cells. The abolishment of luminal cell differentiation in turn prevented the alveologenesis thereby blocking complex TDLUs formation, thus phenocopying DDR1i.

### DDR1 and RUNX share a core transcriptional network that controls breast epithelial differentiation

The similarities in the developmental and morphological defects due to the inhibition of either DDR1 or RUNX1 implies the existence of a shared transcriptional program regulating breast epithelial differentiation. We conducted RNA-Seq analysis on human breast epithelial cells exposed to either collagen alone or collagen in combination with DDR1 or RUNX inhibitors to determine whether there exists a core gene set governed through the activation of these two proteins. Indeed, we found a substantial overlap in differentially expressed genes between DDR1i and RUNXi cells. Over half the genes (316 out of 576) that exhibited differential expression upon DDR1 inhibition overlapped with those affected by RUNX inhibition (Fig. 5A). Moreover, unsupervised clustering analysis of these 316 differentially expressed genes indicated that cells treated with DDR1i, and cells treated with RUNXi closely clustered together, contrasting with the control cells (Fig. 5B). Notably, GSEA of the genes within this shared set reveals that they are implicated in known RUNX1 processes such as tissue development (Coffman, 2003), epithelial-mesenchymal transition (EMT) (Khawaled and Aqeilan, 2017; Lu et al., 2020a), estrogen response (Stender et al., 2010), MYC signaling (Choi et al., 2017), as well as numerous other functions (Supplemental Fig. 4) (Mootha et al., 2003; Subramanian et al., 2005).

**Figure 5:**
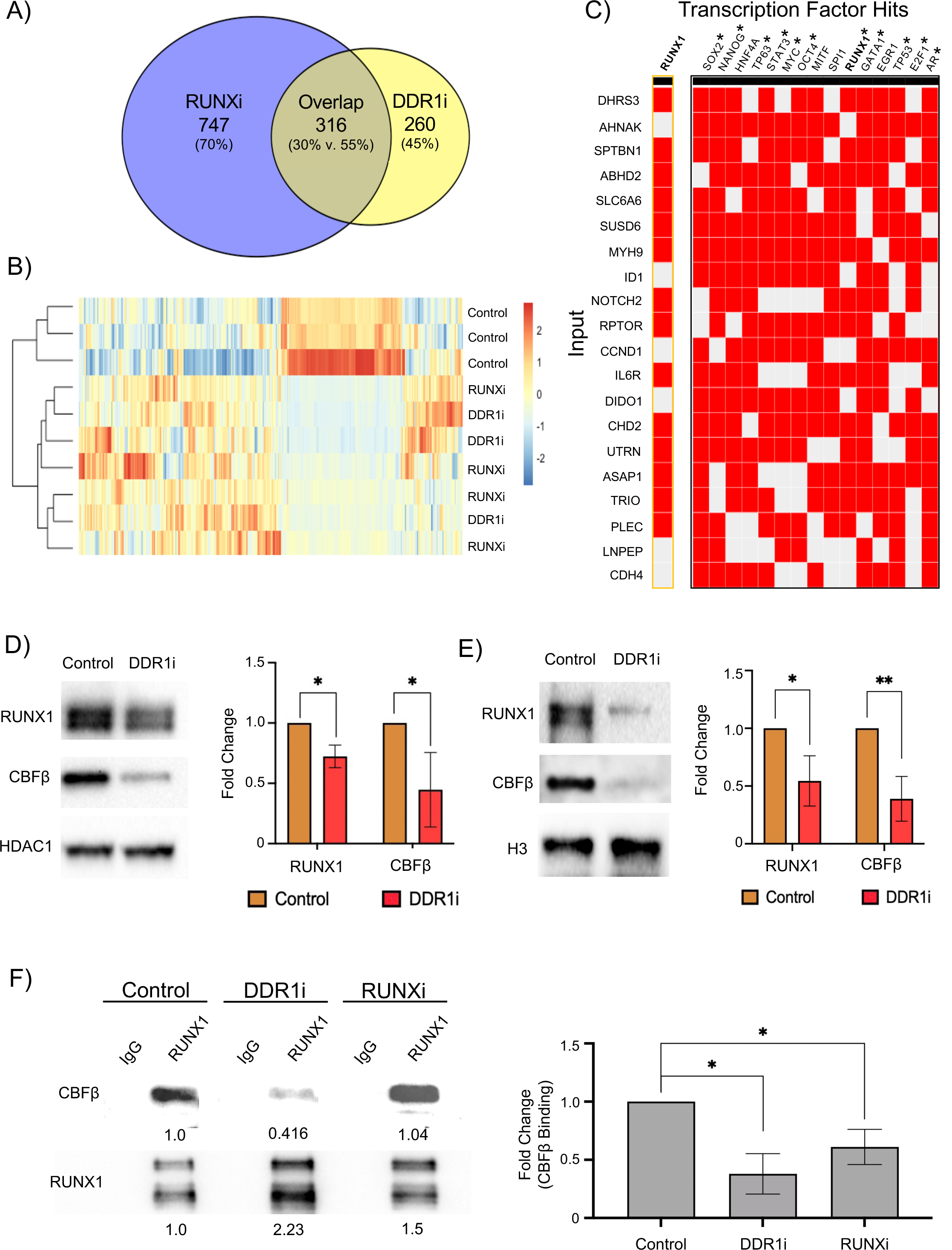
Molecular and transcriptional interactions between DDR1 and RUNX1. A) Venn diagram depicting the overlapping differentially expressed genes from bulk RNAseq data obtained from MCF10A cells treated with Col-1 and DDR1 inhibitor (DDR1i) or Col-1 and RUNX inhibitor (RUNX1i), compared to control. B) Heatmap of the overlapping differentially expressed genes, as compared to the control. C) Hierarchically clustered heatmap visualization of the associations between DDR1-RUNX1 overlapping gene set (input) and Enriched transcription factors, utilizing data from the ChEA database(Lachmann et al., 2010). * Represents transcription factor hits of interest. D) Representative western blot analysis and quantification of RUNX1 and DDR1 expression from nuclear lysates of MCF10A cells treated with collagen and DDR1i (n=3). Data presented as Mean ± SD. E) Representative western blot and quantification of nuclear protein from fractionated lysate obtained from primary patient samples cultured in 3D hydrogels (n=3). Data expressed as Mean ± SD. F) Co-IP of RUNX1 and subsequent blotting and quantification of CBFβ from the lysate of MCF10A samples cultured in 2D treated with collagen and DDR1i. Quantification is derived from n=3 independent experiments and is presented as Mean ± SD. Statistical significance was determined through multiple t-tests, with significance levels indicated as follows: *p-value < 0.05, **p-value < 0.01, ***p-value < 0.001, ****p-value < 0.0001.

We used the ChEA Chip-seq database (Lachmann et al., 2010) visualized with Harmonizome 3.0 (Rouillard et al., 2016) (Fig. 5C) and the TRRUST transcription factor regulator interaction database (Han et al., 2015) visualized with ENRICHR (Xie et al., 2021) (Supplemental Fig. 5) to gain deeper insights into the potential cellular processes governed by the shared 316 DDR1/RUNX target genes. In doing so we found several transcription factors also known to regulate the same target genes. Interestingly, several of these transcription factors also have roles in the regulation of pluripotent stem cell functions, including SOX2, NANOG, and OCT4 (Fig. 5C, Supplemental Fig. 5). TP63, STAT3, MYC, GATA1, TP53, E2F1, and AR, were among other transcription factors (TFs) that also share the overlapping targets of DDR1/RUNX1(Fig. 5C). Notably, these are key TFs that regulate stemness and differentiation programs, in the breast but also male and female reproductive tissues (Chang et al., 2013; Hu et al., 2011; Huang et al.; Jorgez et al., 2021; Murashima et al., 2015; Xu et al., 2023) stratified epithelial tissues (Barbieri and Pietenpol, 2006; Craig et al., 2023; Fang et al., 2020; Portal et al., 2022; Wu et al., 2003), and the hematopoietic system (Delgado and León, 2010; Huang et al., 2014; Kortylewski et al., 2005; Pant et al., 2012; Shimizu and Yamamoto, 2023; Trikha et al., 2011). Input of these transcription factor hits into STRINGDB, a database and visualizer of known protein-protein interactions (Szklarczyk et al., 2022) shows that these proteins form a closely related network, with each of these proteins only being 1-2 interactions away from each other (Supplemental Fig. 6). Collectively, these findings suggest that the DDR1/RUNX1 axis plays a central role in regulating the shift of genes between stem cell state to differentiated states.

As DDR1 is a receptor tyrosine kinase, its ability to regulate RUNX1 transcriptome may also be at the protein level, as well as transcriptional. Indeed, analysis of protein from MCF10A cells stimulated with collagen and DDR1i showed a significant reduction in the total levels of RUNX1 protein as well as its co-factor CBFβ (Fig. 5D). The RUNX1-CBFβ complex is important for RUNX1 protein stabilization and transcriptional activity (Qin et al., 2015, Martinez et al., 2016). Similarly, primary cells grown in 3D and treated with DDR1 inhibitors also exhibited this loss of RUNX1 and CBFβ (Fig. 5E). These results imply that DDR1 inhibition plays a role in the expression of RUNX1 and CBFβ.

As RUNX1-CBFβ complex formation is important to both stability of RUNX1 protein (Qin et al., 2015) and its ability to function as a transcription factor (Tahirov et al., 2001), we looked to see if this interaction was affected by DDR1i. Indeed, co-Immunoprecipitation of CBFβ with RUNX1 revealed a marked reduction in its association with CBFβ in the presence of DDR1i (Fig. 5F). Use of the pan-RUNX inhibitor AI-10-104, a small molecule that inhibits the complex formed between RUNX1 and CBFβ (Illendula et al., 2016), as a positive control also showed a reduction in the association between CBFβ and RUNX1 in treated samples (Fig. 5F). Collectively, this data suggests a core set of stem cell and differentiation genes, controlled by DDR1 through regulation of RUNX1’s expression and its interaction with CBFβ.

### DDR1/RUNX1 axis mutations commonly occur in breast cancer

Breast cancer is fundamentally a disease of misregulated development and differentiation—breast cells no longer properly respond to developmental cues and begin to break free from their lineage-restricted behaviors. Breast cancer progression has consequently been linked to the process of de-differentiation. Thus, factors that control differentiation are often among the most frequently mutated genes in breast cancers. Utilizing the extensive breast cancer database on cBioPortal (de Bruijn et al., 2023; Cerami1 et al., 2012; Gao et al., 2013), in-depth analysis of genetic alteration types by gene demonstrated that DDR1 alterations in breast cancer were predominantly amplifications, RUNX1 alterations mostly consisted of a combination of mutations and amplifications, and CBFβ alterations were mostly mutations and deep deletions (Fig. 6A). These mutations occur at rates like those seen with EGFR mutations, a commonly occurring mutation, at frequencies between 3∼5% of all breast tumors (Fig. 6A). Further analysis of this database revealed a highly significant tendency for the co-occurrence of mutations in DDR1 and RUNX1 as well as in DDR1 and CBFβ, while RUNX1 and CBFβ mutations tend to be mutually exclusive, as expected (Fig. 6B). Together this data suggests that the DDR1-RUNX1 axis is often perturbed in breast cancer tumors.

**Figure 6:**
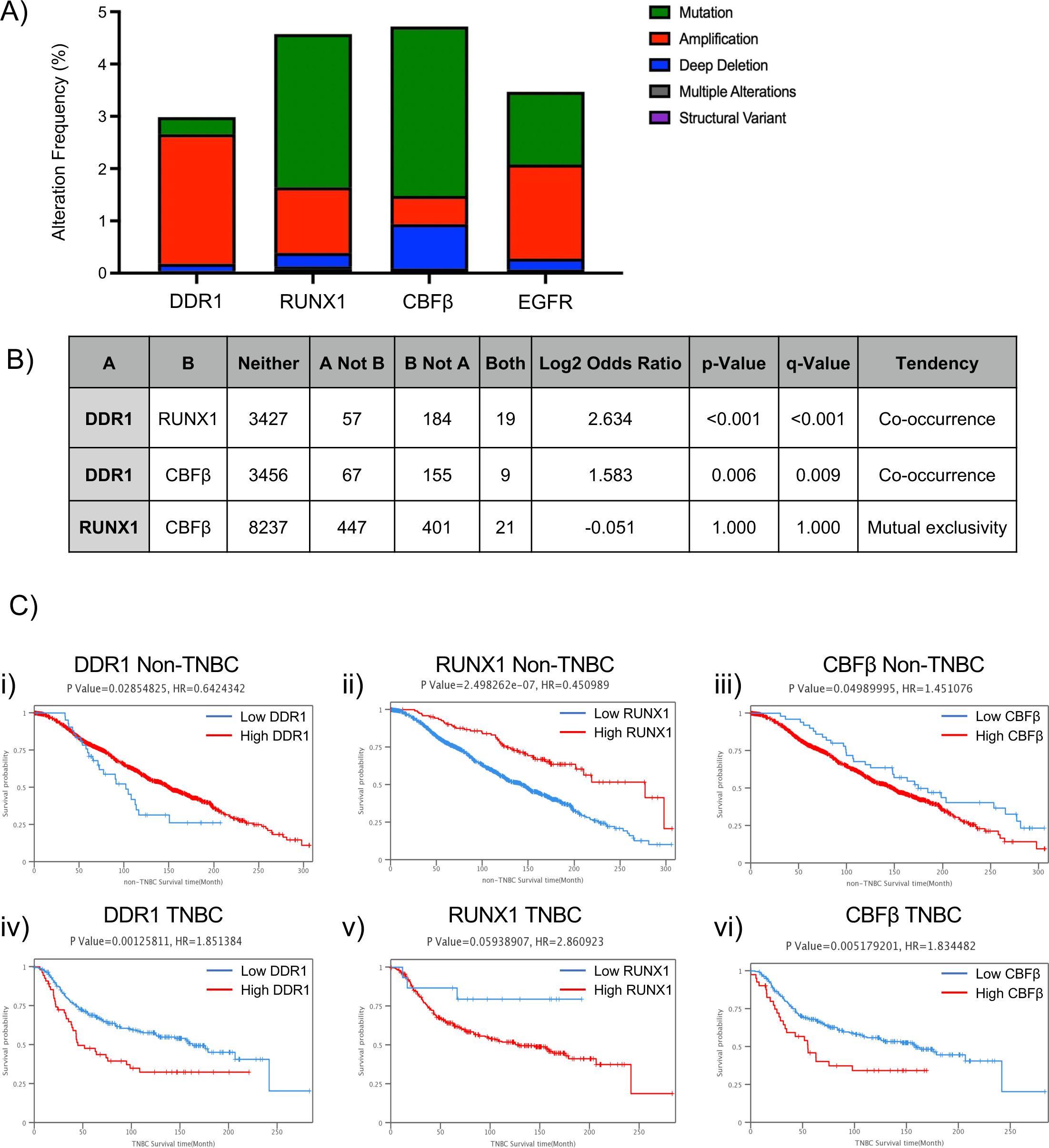
Comprehensive analysis of gene expression and clinical outcomes. A) Genomic alteration frequencies of RUNX1, DDR1, CBFb, and EGFR in human breast cancer. Mutational frequency from 10,363 breast cancer (BC) samples in 9,776 patients, categorized by mutation type, of genes of interest and well-known regulators in BC from the cBioPortal database. B) Mutual exclusivity of DDR1, RUNX1, and CBFβ from the mutational data of 10,363 BC samples in 9,776 patients compiled on the cBioPortal database. C) Kaplan-Meier OS curve based on high (red) and low (blue) DDR1 (i), RUNX1 (ii), or CBFβ (iii) in triple negative breast cancer (TNBC) or non-TNBC (iv-vi) from METABRIC data on the Breast Cancer Integrative Platform.

To assess the clinical relevance of DDR1, RUNX1, and CBFβ expression in breast cancer, we used METABRIC data from the Breast Cancer Integrative Platform (Wu et al., 2017) to produce Kaplan-Meier curves stratifying the effects of their expression by breast cancer subtype. The results unveiled a discernible pattern: elevated expression of both DDR1 and RUNX1 had a positive influence on the prognosis of non-triple negative breast cancers (ER^+^/PR^+^/HER2^+^) (Fig. 6C, panels i-ii), while increased expression of CBFβ had a slight negative prognosis in these cancer types (Fig. 6C, panel iii). In stark contrast, heightened expression of all three genes was associated with a negative prognosis in TNBC. (Fig. 6C, panels iv-vi). These findings underscore a connection between the expression patterns of DDR1 and RUNX1 across different cancer types and their involvement in stem cell states and lineage differentiation, that could have relevant clinical implications for treatments in different breast cancer subtypes.

## Discussion

The presented data establishes an association between the activation of DDR1 and the transcriptional activity of during breast tissue development and maturation. Our findings connect previous observations about these two proteins (Hong et al., 2017; Rauner et al., 2021; Sokol et al., 2015; Suh and Han, 2011) and indicate that they jointly play pivotal roles during the early phases of breast organogenesis, encompassing induction and patterning processes. Failure to activate DDR1 and RUNX1 during these developmental stages impedes both cellular lineage proliferation and the formation of alveolar structures. Furthermore, we observe that DDR1 inhibition reduces the interaction between RUNX1 and CBFβ. Given CBFβ’s canonical role in enhancing the affinity between RUNX proteins and DNA (Malik et al., 2019; Ogawa et al., 1993; Rose et al., 2020) and considering the predominant expression of RUNX1 in breast tissue(Mercado-Matos et al., 2017), this disruption in binding exerts substantial effects on the RUNX1 transcriptome. Notably, 55% of the differentially expressed genes in breast cells subjected to DDR1 inhibition exhibit changes consistent with those observed in cells exposed to a RUNX1-CBFβ binding inhibitor. According to the ChEA database (Lachmann et al., 2010), (Rouillard et al., 2016), a significant proportion (54%) of these differentially expressed genes are recognized as direct RUNX1 target genes, while the remainder may be attributed to transcription factors whose expression is regulated by RUNX1. While we were able to see a clear connection between DDR1 and RUNX1 through modulation to both expression and activity with inhibition, we were not able to determine if this modulation was the consequence of a direct or indirect interaction between the two proteins. Further work needs to be done to determine how the RUNX1-CBFβ complex is regulated downstream of DDR1 activation.

DDR1’s control over RUNX1 seems to start through impacting RUNX1 and CBFβ’s ability to interact with one another. Preventing this interaction could trigger a transcriptional autoregulatory cycle and create a redundant signal leading to not only the lack of complex formation but also a decrease in RUNX1 mRNA and protein expression, pointing to the importance of control over this core gene set and the relationship of these proteins. The RUNX1 and CBFβ complex have been previously observed to affect both the protein stability of RUNX1 (Qin et al., 2015) and has the ability to promote RUNX1 transcription (Martinez et al., 2016), both of which is supported by the loss of nuclear RUNX1 protein and mRNA as seen in our data. There is also limited literature on the control over the localization of the RUNX1-CBFβ complex; however, data suggests that it is necessary for this complex to form in the cytoplasm prior to its trafficking into the nucleus (Malik et al., 2019). Further work should be done to investigate if the alterations to their interaction, caused by DDR1 inhibition, also cause changes in the trafficking of this complex. Driven by DDR1, these alterations to RUNX1 ensure swift regulation of the RUNX1 transcriptome by initiating redundancies of control at both the protein and transcriptional level.

While still requiring further investigation, our data offers valuable insights into the functional roles of RUNX1 in breast tissue. These roles seem to be split by the phase of development and the differentiation state of the cells. During induction, inhibition of the DDR1-RUNX1 axis blocks differentiation and prevents bipotent progenitors and stem cells from giving rise to lineage committed progenitors. During patterning disruption of signaling through the DDR1-RUNX1 axis prevents normal physiological morphogenesis. Inside the cell, we see that inhibition of DDR1 leads to loss of RUNX1 mRNA expression within progenitor and basal cell types, resulting in an overall depletion of RUNX1 and its associated binding partner CBFβ proteins. Previous transplant and organoid models have demonstrated that progenitor and basal cells are exposed to the extracellular matrix and are the initial cell types to develop (Arendt et al., 2010, 2014; Russo and Russo, 2004). Moreover, the effects of RUNX1 and DDR1 inhibition manifest exclusively in those early cell types when treatment is initiated during the initial stages of breast development, indicating an influence of these proteins on stem cells and early progenitors.

Pathway enrichment analysis tools showed that the overlapping differentially expressed genes upon DDR1i and RUNXi are under the regulatory influence of stem and differentiation-related transcription factors, including pluripotency inducing SOX2, OCT4, and Nanog (Allouba et al., 2015; Luo et al., 2013; Shi and Jin, 2010; Zhang and Cui, 2014). These three transcription factors, along with E2F1 (Guy et al., 1996), are recognized for their ability to repress differentiation, whereas other transcription factor hits known to regulate these genes, such as TP53, STAT3, EGR1, and GATA1, promote differentiation (Briegel et al., 1996; Guerquin et al., 2013; Lu et al., 2020b; Sivakumar et al., 2022). Additionally, the remaining transcription factors on the list, including AR, MYC, and TP63, are known to exert influences on both stem and differentiated states(Huang et al., 2014; Melnik et al., 2019; Nekulova et al., 2011).

These transcription factors can be categorized into two distinct groups: inducers of stem programs that function as proto-oncogenes (SOX2, OCT4, Nanog, E2F1, MYC) (Dang, 2012; Narayan et al., 2017; Noh et al., 2012; Schaefer and Lengerke, 2020; Yang and Sladek, 2018), and drivers of differentiation that function as tumor suppressors (TP53, STAT3, EGR1, and GATA1) (Aigner et al., 2019; Aubrey et al., 2016; Wang et al., 2021; Zheng and Blobel, 2010). This categorization serves to clarify the dual functionality of RUNX1 expression observed in development and across various breast cancer subtypes. Notably, it has been observed that RUNX1 is interconnected with these transcription factor hits either directly or through a known interaction with another protein known to be involved in the regulation of this core DDR1/RUNX1 gene set (Supplemental Fig. 4). The regulatory interactions between RUNX1 and these proteins are often seen to be on a transcriptional level (Choi et al., 2017; Elagib et al., 2003; Fernández et al., 2023; Lachmann et al., 2010; Masse et al., 2012), promoting vast changes in transcriptional programming. This extensive network of interactions underscores the pivotal role of RUNX1 activation in orchestrating the transcriptional activity necessary for determining cell fate.

The range of functions associated with the aforementioned transcription factor hits delineates the seemingly dual role that RUNX1 plays downstream of DDR1, involving both the suppression of stem-related genes and the activation of epithelial ones. A particularly intriguing observation is that nearly 70% of the overlapping gene set (219 out of 316 genes) exhibits upregulation in response to the loss of either RUNX1 or DDR1. This points to RUNX1 playing a predominantly inhibitory role in the breast and, when activated, preventing the transcription of genes contributing to a stem like state, while activating genes required for differentiation.

Recently, DDR1, RUNX1, and CBFβ have all garnered attention as potential clinical targets in breast cancer (Ariffin, 2022; Fowler et al., 2020; Han et al., 2022; Khawaled and Aqeilan, 2017; Malik et al., 2019). Current evidence indicates that RUNX1 and CBFβ are among the 30 most frequently mutated genes in breast cancer (Banerji et al., 2012; Koboldt et al., 2012; Pereira et al., 2016), with a prevalence of approximately 4-5% in breast cancer cases (Pereira et al., 2016). While DDR1 mutations in breast cancer are estimated to occur at a rate of 2-4% each, it is worth noting that these figures may be underreported as many genomic studies on breast cancer have not explicitly explored DDR1. It is also worth noting that CBFβ mutations may also be under reported as the loss of the CBFβ’s chromosomal loci at 16q22 represents one of the most frequent and earliest genomic alterations observed in breast cancer, affecting roughly 50% of all cases (Cleton-Jansen et al., 2000; Przybytkowski et al., 2014).

Human tumor sequencing data reveals a substantial co-occurrence of mutations between DDR1 and either RUNX1 or CBFβ, whereas such co-occurrence is not necessarily observed between RUNX1 and CBFβ. This data is in line with our findings, as mutations to DDR1 and either RUNX1 or CBFβ would have two distinct effects, while loss of both RUNX1 and CBFβ would be largely redundant. In these cases, concurrent loss mutations of DDR1 and a member of the RUNX1-CBFβ complex could allow for survival advantages by manipulating their role in the balance of stem and differentiated states, or by promoting their normal function of proliferation. Further examination of this co-occurring mutation across all cancer studies on cBioPortal showed that this mutational relationship between DDR1 and either RUNX1 or CBFβ remained strongly significant, indicating that this axis is worth investigating outside of breast cancer. Given the development of potent and selective inhibitors targeting both RUNX1 and DDR1 for potential clinical therapies against various localized tumors, research aimed at deepening our understanding of the role of the DDR1/RUNX1 axis in oncogenesis and normal development.

## Conclusions

Our investigation employing a cutting-edge breast organoid model generated in a 3D hydrogel has shed light on a previously unrecognized yet crucial player within the DDR1 signaling pathway—RUNX1. The activation of DDR1, triggered by the binding of its ligand, collagen, controls the association of RUNX1 with CBFβ, and thus their downstream transcriptional processes. This newly uncovered DDR1-RUNX1 axis operates as a potent stem cell transcription factor signaling node, orchestrating differentiation, and significantly impacting the morphological characteristics of breast epithelial structures. The clinical implications of these proteins’ expression in cancer are breast cancer subtype specific, warranting further scrutiny to harness the full potential of inhibitor-based therapeutic interventions.

## Experimental Procedures

### Resource Availability

**Corresponding Author**

Information requests can be directed to corresponding author Charlotte Kuperwasser

### Ethics Statement

Primary tissues that normally would have been discarded as medical waste post-surgery were obtained in compliance with all relevant laws, using protocols approved by the institutional review board at Maine Medical Center and Tufts Medical Center. All tissues were anonymized before transfer to prevent tracing back to specific patients; for this reason, this research was provided exemption status by the Committee on the Use of Humans as Experimental Subjects at the Massachusetts Institute of Technology, and at Tufts University Health Sciences (IRB# 13521). All enrolled patients in this study signed an informed consent form agreeing to participate in this study and for publication of the results.

### 2D Cell Culture of MCF10 Cells

MCF10A (ATCC CRL-10317) and MCF10F (ATCC CRL) cells were cultured in DMEM/F12 (Corning) supplemented with 10 μg/mL insulin (Sigma), 20 ng/mL hEGF (E9644, Sigma Aldrich), 500 ng/mL hydrocortisone (Sigma), 100 ng/mL cholera toxin (Sigma-Aldrich), 5% Horse Serum (Gibco), and 1x Antibiotic/antimitotic (Corning).

### Primary Sample Preparation

Primary tissue samples from reduction mammoplasties of healthy women were dissociated through a 3 mg/mL collagenase (Roche Life Science) solution and sorted via density gradient. Epithelial clusters were dissociated to single cells using 0.25% trypsin-EDTA (Gibco) and filtered through a 40 mm mesh filter after fibroblast removal.

### Collagen Stimulation Assay

MCF10A and MCF10F cells were seeded at densities of 1e6 cells for 10 cm plates or 2.5e6 cells for 15 cm plates. Cells were treated as control or with 2 µM DDR1 inhibitor DDR1-in-1 (Tocris 5077) or 5 µM RUNX1 inhibitor AI-10-104 (Aobious AOB17076). A solution of media, collagen (0.05 mg/mL), and 0.1 N NaOH for polymerization was placed on cells during media changes. After 24 hours, collagen was removed, cells were lifted with 0.25% trypsin-EDTA, and prepared for downstream assays.

### 3D Hydrogel Model Culture

Single cell primary tissue samples and MCF10A cells were mixed in a suspension of 1.7 mg/mL rat tail collagen I (Corning), 40 μg/mL laminin (Thermo Fisher Scientific), 20 μg/mL fibronectin (Gibco), and 10 μg/mL hyaluronic acid (Sigma), adjusted to pH 7.3 with 0.1 N NaOH. Hydrogels were plated in a four-chamber slide (Falcon) as a mold, polymerized for 1 h at 37 °C, and overlaid with MEGM medium (Lonza CC-3150) supplemented with 1x Antibiotic/Antimitotic, 1x Glutamax (Gibco). Structures were dissociated with collagenase and trypsin-EDTA, depending on downstream assays.

### Inhibitor Time course

Single cell primary tissue samples were seeded into gels in three conditions: control, chronic DDR1 or RUNX inhibitor treatment starting day 0 (induction), and chronic DDR1 or RUNX inhibition starting day 7 (patterning). Samples were matured for up to 28 days and scored blinded for structure types.

### RNA Isolation and Quantitative RT-PCR

Cells were pelleted, and RNA was isolated using the RNAeasy kit (Qiagen). cDNA was produced with the iScript cDNA kit (Bio-Rad), and qRT-PCR was performed with Sybr green (Bio-Rad). Primers for target genes are listed below.

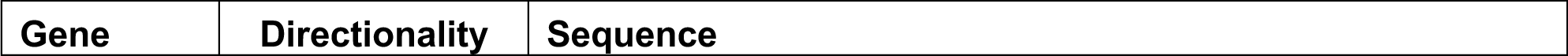
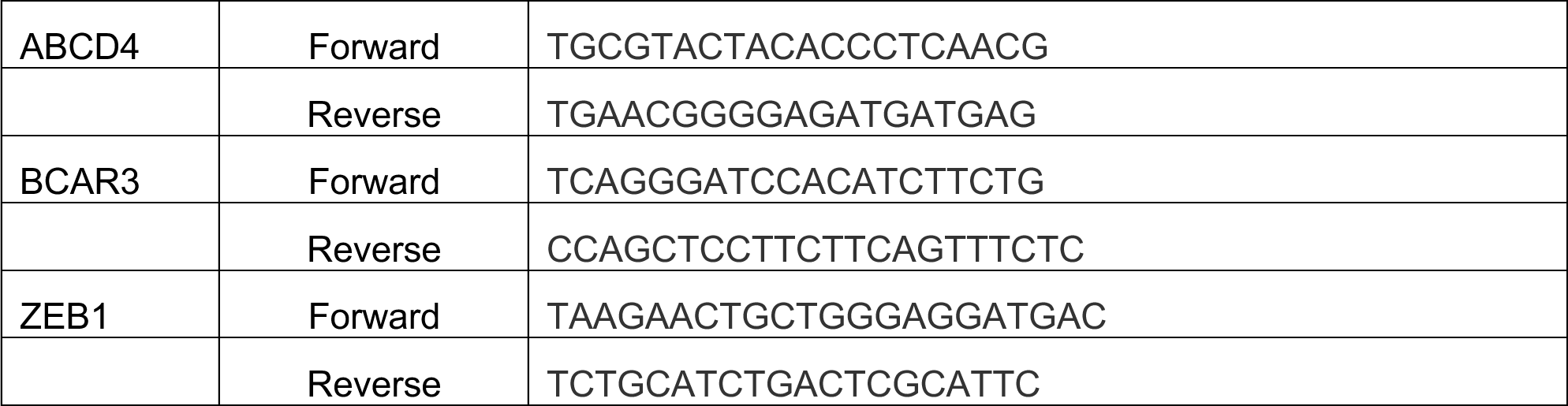

### SC-RNA Seq

Previously published scRNA-seq data from organoids(Rauner et al., 2021). scRNA-seq data were analyzed using Seurat v3 (Stuart et al., 2019) for data integration, normalization, and feature selection. Briefly, raw data was loaded and integrated into one Seurat object using the merge function. Filtering removed cells with < 200 or >2500 genes and mitochondrial content greater than 7.5%. Genes detected in less than 3 cells were dropped from analysis. The data was normalized by multiplying transcripts by a factor of 10,000 followed by log-transforming the data. Variable features used for analysis were identified by using the FindVariableFeatures function, with a low cutoff of 0.0125 and a high cutoff of 5 for dispersion and a low cutoff of 0.1 and a high cutoff of 0.8 for average expression. The data was integrated by the FindIntegrationAnchors and IntegrateData functions, which identify the anchors to integrate the two datasets, and then integrates them together. Cells were then clustered using K-nearest neighbor (KNN) graphs and the Louvain algorithm using the first 10 dimensions from principal component analysis. Clustered cells were visualized by tSNE embedding using the default settings in Seurat. Clusters were called using the FindClusters function with a resolution of 1. To identify differentially expressed genes between cell clusters, we utilized the FindAllMarkers function to identify features detected in >10% of a cell cluster compared to all other cells. Pathway analysis to identify enriched biological pathways associated with differentially expressed genes was done using established databases, such as PanglaoDB (Franzén et al., 2019). The top 15 differentially expressed markers were used to determine gene expression location.

### Western Blot

Cells grown in 2D or isolated from 3D hydrogels were pelleted by centrifugation and were fractionated using the NE-PER Nuclear and Cytoplasmic Extraction Reagents (Thermo, 78833), containing both 1x protease inhibitor cocktail (Cell Signaling Technology) and 1x phosphatase inhibitor (Cell Signaling Technology) according to manufacturer’s protocol. Nuclear fraction samples were separated via NuPAGE gel (Invitrogen) and transferred to PDVF (BioRad) to be blocked in 5% BSA (Rockland). Blots were then incubated with primary antibody overnight at 4°C and with secondary antibody for 1 hour at room temperature. Immunoblot membranes were developed using a chemiluminescent substrate (Thermo Fisher Scientific) and imaged with the Chemidoc XRS+ with Image Lab 6.0.1 software (BioRad, Hercules, CA). ImageJ2 (Version 2.8.0/1.53t) was used to densitometry quantifications. Primary Antibodies used were: RUNX1 (4336, Cell Signaling Technology, Clone D33G6, 1:1000), CBFβ (A303-549A, Bethyl Laboratories, 1:1000), HDAC1 (5356, Cell Signaling Technology, Clone 10E2, 1:1000), H3 (9715, Cell Signaling Technology, 1:1000). Secondary antibodies used were Goat anti-Rabbit (7074, Cell Signaling Technology, 1:1000) and Goat anti-Mouse (7076, Cell Signaling Technology, 1:1000).

### Co-Immunoprecipitation

MCF10A cells from 2D collagen stimulation assays were pelleted by centrifugation and lysed using 1X RIPA Buffer containing both 1x protease inhibitor cocktail and 1x phosphatase inhibitor Lysate was precleared with Protein-A Magnetic Beads (73778, Cell Signaling Technology).

Cells were then incubated in immunoprecipitive antibody overnight at 4°C. Antibodies and attached proteins were conjugated to the magnetic beads at room temperature for 40 minutes. Samples were separated via SDS-PAGE gel and transferred to PDVF to be blocked in 5% BSA.

Blots were then incubated with primary antibody overnight at 4°C and with secondary antibody for 1 hour at room temperature. Immunoprecipitative antibody was RUNX1 (HPA004176, SIGMA, 5ug/mg lysate). Primary antibodies were RUNX1 Ms (sc-365644, Santa Cruz, A-2, 1:1000), CBFβ Ms (67885-1, ProteinTech,1D7F2, 1:1000), and secondary antibody: Goat anti-Mouse (7076, Cell Signaling Technology, 1:1000).

### Microscopy

Immunofluorescence images captured using Nikon AXR (Nikon Microscopy). Brightfield images captured using Nikon Eclipse Ti-U (Nikon Microscopy), using SPOT 5.6 software.

### Immunofluorescence

Cells and hydrogels were fixed with 4% Paraformaldehyde (Fisher Scientific), permeated with 0.1% Triton 100X, and incubated at 4°C for 18 hours with primary antibodies: E-Cad (13-1700, Thermo Fisher, HECD-1, 1:100) and CK-14 (RB-9020, Thermo Fisher, 1:300). Samples were then incubated at 4°C for 18 hours with secondary antibodies: DAPI (D1306, Life Technologies, 1:1000), AF488 (A11008, Invitrogen,1:1000), AF555 (A21424, Invitrogen,1:1000), Phalloidin-AF647 (A22289, Invitrogen,1:500).

### Live Imaging

Primary single cells, isolated as described above, were incubated with the cell tracking dye Cytopainter Green (1:500, cat# ab138891, Abcam) for 30 minutes and then were washed and seeded at a concentration of 100 cells per 20 uL hydrogel. For gel fabrication, 20 uL hydrogel drops were deposited onto the center wells of a 96-well plate (Corning, #3603). Gels were allowed to incubate for one hour at 37 °C until fully polymerized. 80uL of MEGM was then added to each well and gels were gently lifted off the well surface with a pipette tip. Cultures were immediately placed in a pre-warmed incubator chamber (Okolab Inc) enclosed over a Nikon Eclipse Ti2-AX confocal microscope (Nikon Microscopy). Images of selected points were collected starting immediately after the addition of media, and every 30-45 minutes after, in both brightfield and with A488 laser at 4x magnification and 2.5x zoom across nine z-positions. 20-40uL of MEGM was added to the culture twice a week to maintain proper growth factors and liquid volume to prevent hydrogels from drying out. Cultures were live imaged for 18-21 days. Analysis and production of videos across locations and timepoints was performed using NIS-Elements (Nikon) and Premiere Pro (Adobe) software.

### Flow Cytometry

Patient sample derived structures, either control or inhibited during patterning at Day 7, were dissociated and pelleted by centrifugation. Samples were washed and stained with the antibodies CD49f-FITC (555736, BD Biosciences, GoH3, 1:20) and EPCAM-PE (347198, BD Biosciences, 1:20). Samples were run on LSRII. FlowJo (Version 10.9.0) was used for visualization and quantification.

### RNA-Seq

mRNA isolated from MCF10A cells in a collagen stimulation assay was run on the Illumina NextSeq 6000.

### RNA-Seq Analysis

Read Alignment to the human genome was performed using STAR (Dobin et al., 2013) with the CRCh37/hg19 assembly. Library normalization as well as differential expression testing was performed using R with DESeq2 (3.17) (Love et al., 2014). Sample SSR107 was removed as it was deemed outlier by PCA. Differential gene analysis was conducted with significance determined by log fold change greater than or less than 0 with a p value of less than 0.05. Gene Ontological analysis was performed using the ChEA dataset (Lachmann et al., 2010) on Harmonize (Version 3.0) and with the TRRUST(Han et al., 2015) dataset from Enrichr (Xie et al., 2021). Heat maps and Venn Diagrams were created in R using Pheatmap (Kolde, 2023) and ggVennDiagram (Gao et al., 2021) respectively.

### Mutational Analysis

RUNX1, CBFβ, and DDR1 were probed for breast cancer alteration frequency on cBioPortal using primary tumor BC data from 20 studies broken down by cancer type (Cerami1 et al., 2012). Mutational status across breast cancer subtype by PAM50 and exploration into mutational mutual exclusivity were also conducted using these studies on cBioPortal. Kaplan Meier survival curves were produced using METABRIC data the Breast Cancer Integrative Platform (BCIP), plotting overall survival based upon transcriptome analysis of either a triple negative status, or non-triple negative (i.e., the expression of at least one receptor ER/PR/HER2).

### Statistics

All statistics were performed using GraphPad Prism 8-10. Student’s t-tests (two sided) were performed as a determinant of significance unless otherwise stated. Data expressed as Mean ± SD. Significance levels indicated as follows: * p-value < 0.05, ** p-value < 0.01, *** p-value < 0.001, **** p-value < 0.0001.

## Acknowledgements

We gratefully acknowledge Albert Tai, Irena Grinvald, and Michael Berne at the Tufts Genomics core for high-throughput sequencing services. Stephen Kwok and Allen Parmelee at Tufts Flow Facilities for flow cytometry support. Karla Murga, Daniela Requena, and Megan Maloney at Tufts Biomedical Repository for Tissue Support. This research was supported by the following: NIGMS (7R01GM124491, CK), Breast Cancer Research Foundation (CK) and FTC Breast Cancer Foundation (CK)

## Author Contributions

CJT, GR, NCT, MEP, DF, YM and CK conceived the project and designed experiments. CJT performed experiments. CJT and NCT performed sequencing analysis. CJT and CK wrote the manuscript.

---

**Supplemental Figure 1: Effects of DDR1i on breast organoid development**

A) Quantification of the total number of organoids that formed following DDR1i treatment during induction. Data presented as Mean ± SD (n= 4 gels/primary patient samples). B) Quantification of the total number of organoids that formed following DDR1i treatment during Patterning. Data presented as Mean ± SD (n= 4 gels/primary patient samples).

**Supplemental Figure 2: Differential expression analysis in response to DDR1 inhibition.**

A) Violin plots showing the distribution of DDR1 expression across epithelial breast cell types. B-I) Violin plots showing the distribution of expression in RUNX1 target genes: ANP32B, PLEC, ID1, STAT3, and the expression of related transcription factors RUNX2, RUNX3 and CBFβ across epithelial clusters in primary tissue organoids grown for 14 days under control DDR1i or DDR1r conditions. J) Quantification of RUNX1 target gene expression from three different patient samples (i-iii). Data expressed as Mean ± SD. K) Quantification of RUNX1 target gene expression from MCF10F cells in a 2D collagen stimulation assay. Data expressed as Mean ± SD. Statistical significance was determined through multiple t-tests, with significance levels indicated as follows: *p-value < 0.05, **p-value < 0.01, ***p-value < 0.001, ****p-value < 0.0001.

**Supplemental Figure 3: Effects of RUNXi on breast organoid development**

A) Quantification of the total number of organoids that formed following RUNXi treatment during induction. Data presented as Mean ± SD (n= 4 gels/primary patient samples). B) Representative brightfield images depicting primary patient samples cultured in a 3D environment, comparing control conditions with DDR1 and RUNX inhibitor-treated conditions initiated during patterning. Scale bar = 200 μm. C) Quantification of the total number of organoids that formed following RUNXi treatment during Patterning. Data presented as Mean ± SD (n= 4 gels/primary patient samples).

**Supplemental Figure 4: GSEA analysis of DDR1-RUNX1transcriptome**

A). Gene set enrichment analysis (GSEA) interrogating core DDR1-RUNX1 gene set with the top 20 Hallmark Gene sets from Molecular Signatures Database (MSigDB).

**Supplemental Figure 5: Transcription factor insights into core DDR1-RUNX1 gene set**

A) Hierarchically clustered heatmap presenting the association between various transcription factor hits and genes differentially expressed from our overlapping gene set (input), utilizing data from the TRRUST database. Red squares indicate that the associated transcription factor is known to regulate the gene in this dataset. Generated with ENRICHR.

**Supplemental Figure 6: String analyses of transcription factor hits**

A) STRINGDB network of protein-protein interactions between the top transcription factor hits from ChEA database known to regulate the core DDR1 and RUNX1 transcriptome.

**Supplemental Video 1: Generation of miniaturized breast structure from a single cell1**

A) Time-lapse analysis capturing the dynamic process of breast organoid formation from a single cell over an 18-day period.

