## Supplemental Figures for "DDR1 regulates RUNX1-CBFβ to control breast stem cell differentiation"

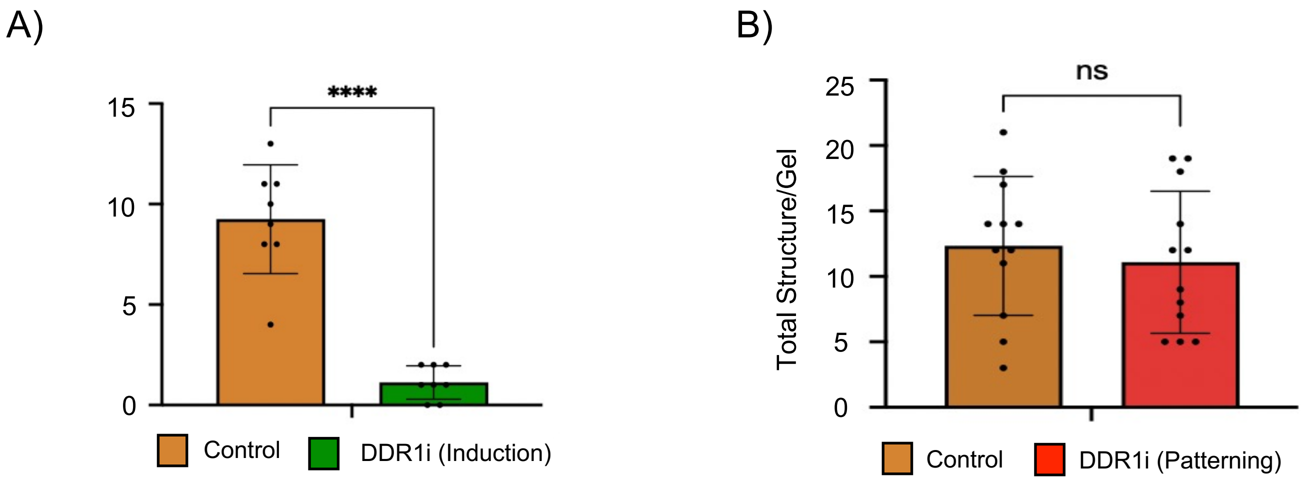


Supplemental Figure 1

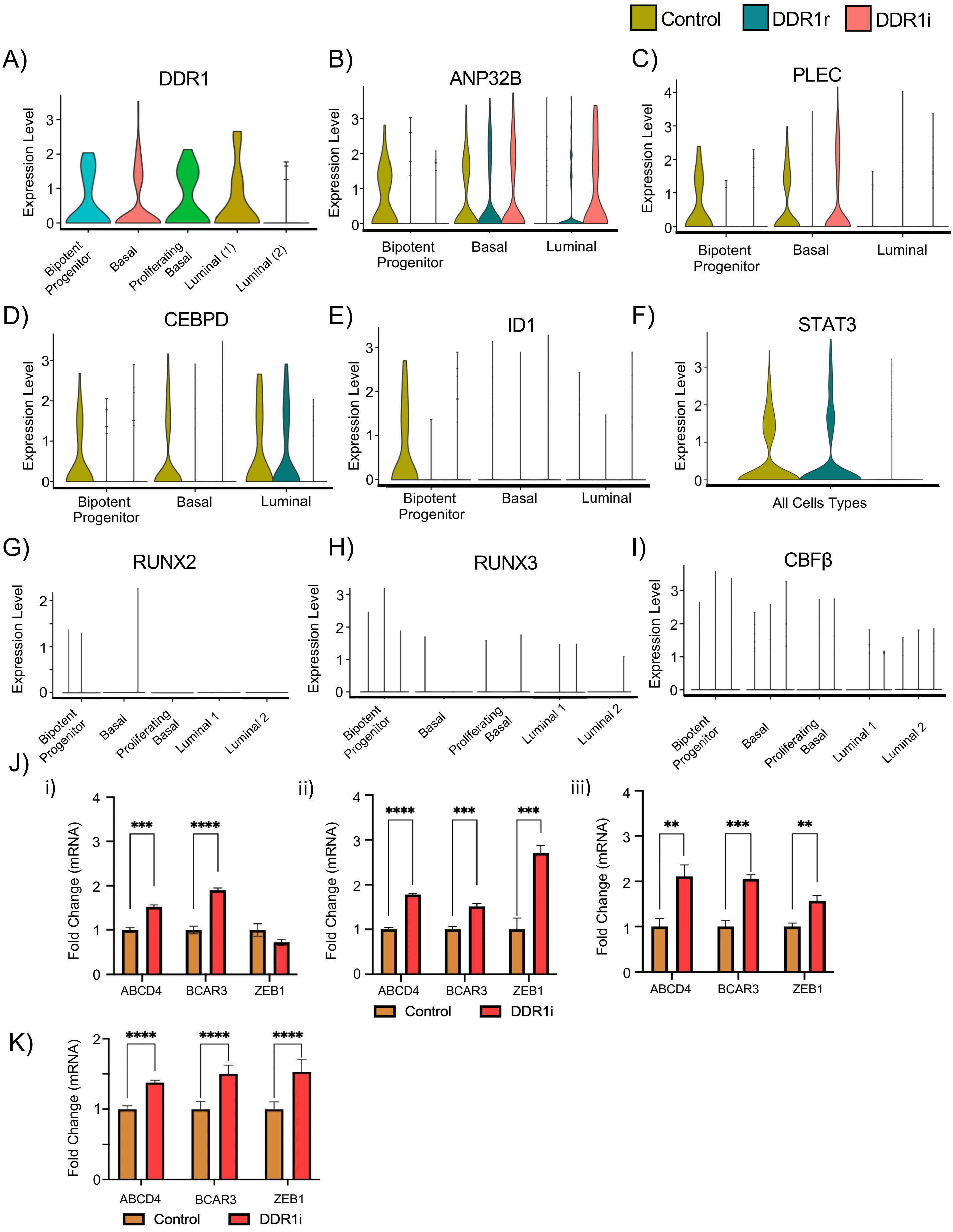


Supplemental Figure 2

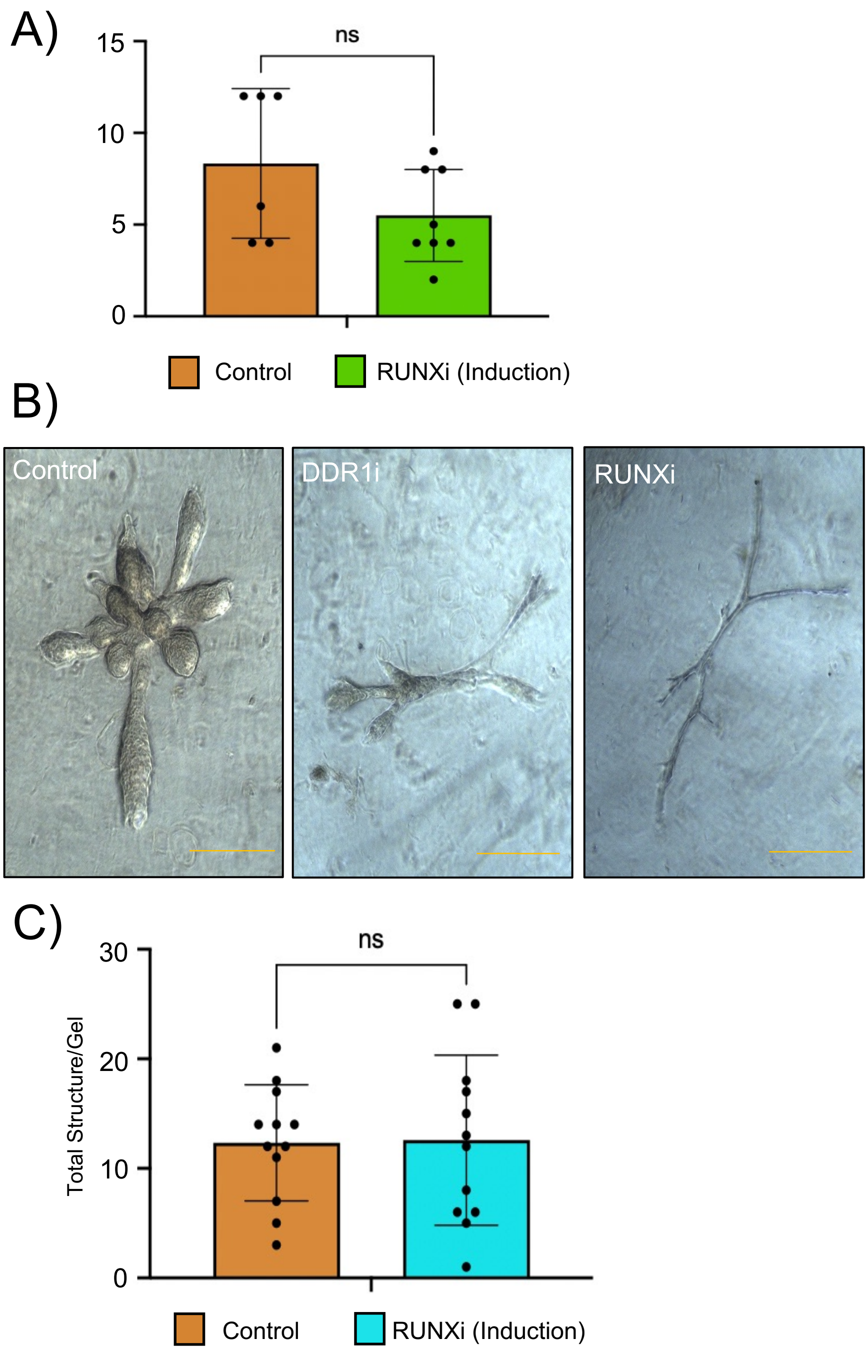


Supplemental Figure 3

A)

| Hallmark Gene Set | # of Bookmarked Genes | p-value | FDRq-value |
|---|---|---|---|
| **Mitotic Spindle** | 21 | $3.03\ e^{-18}$ | $1.51\ e^{-16}$ |
| **Myc Targets** | 18 | $1.34\ e^{-14}$ | $3.36\ e^{-13}$ |
| **Oxidative Phosphorylation** | 14 | $3.48\ e^{-10}$ | $5.81\ e^{-9}$ |
| **Apical Junction** | 11 | $3.21\ e^{-7}$ | $3.21\ e^{-6}$ |
| **Epithelial Mesenchymal Transition** | 11 | 3.21 e-7 | 3.21 e-6 |
| **IL2 Stat5 Signaling** | 10 | $2.5\ e^{-6}$ | $2.09\ e^{-5}$ |
| **E2F Targets** | 9 | $1.93\ e^{-5}$ | $1.07\ e^{-4}$ |
| **Estrogen Response Early** | 9 | $1.93\ e^{-5}$ | $1.07\ e^{-4}$ |
| **Estrogen Response Late** | 9 | $1.93\ e^{-5}$ | $1.07\ e^{-4}$ |
| **NOTCH Signaling** | 4 | $8.75\ e^{-5}$ | $4.38\ e^{-4}$ |
| **Myogenesis** | 8 | $1.27\ e^{-4}$ | $5.77\ e^{-4}$ |
| **TGF Beta Signaling** | 4 | $6.78\ e^{-4}$ | $2.31\ e^{-3}$ |
| **UV Response (Down)** | 6 | $7.17\ e^{-4}$ | $2.31\ e^{-3}$ |
| **G2M Checkpoint** | 7 | $7.4\ e^{-4}$ | $2.31\ e^{-3}$ |
| **MTORC1 Signaling** | 7 | $7.4\ e^{-4}$ | $2.31\ e^{-3}$ |
| **TNFA Signaling vis NFkB** | 7 | $7.4\ e^{-4}$ | $2.31\ e^{-3}$ |
| **Complement** | 6 | $3.77\ e^{-3}$ | $1.11\ e^{-2}$ |
| **Myc Targets V2** | 3 | $9.01\ e^{-3}$ | $2.5\ e^{-2}$ |
| **Heme Metabolism** | 5 | $1.65\ e^{-2}$ | $4.13\ e^{-2}$ |
| **P53 Pathway** | 5 | $1.65\ e^{-2}$ | $4.13\ e^{-2}$ |

Supplemental Figure 4

A)

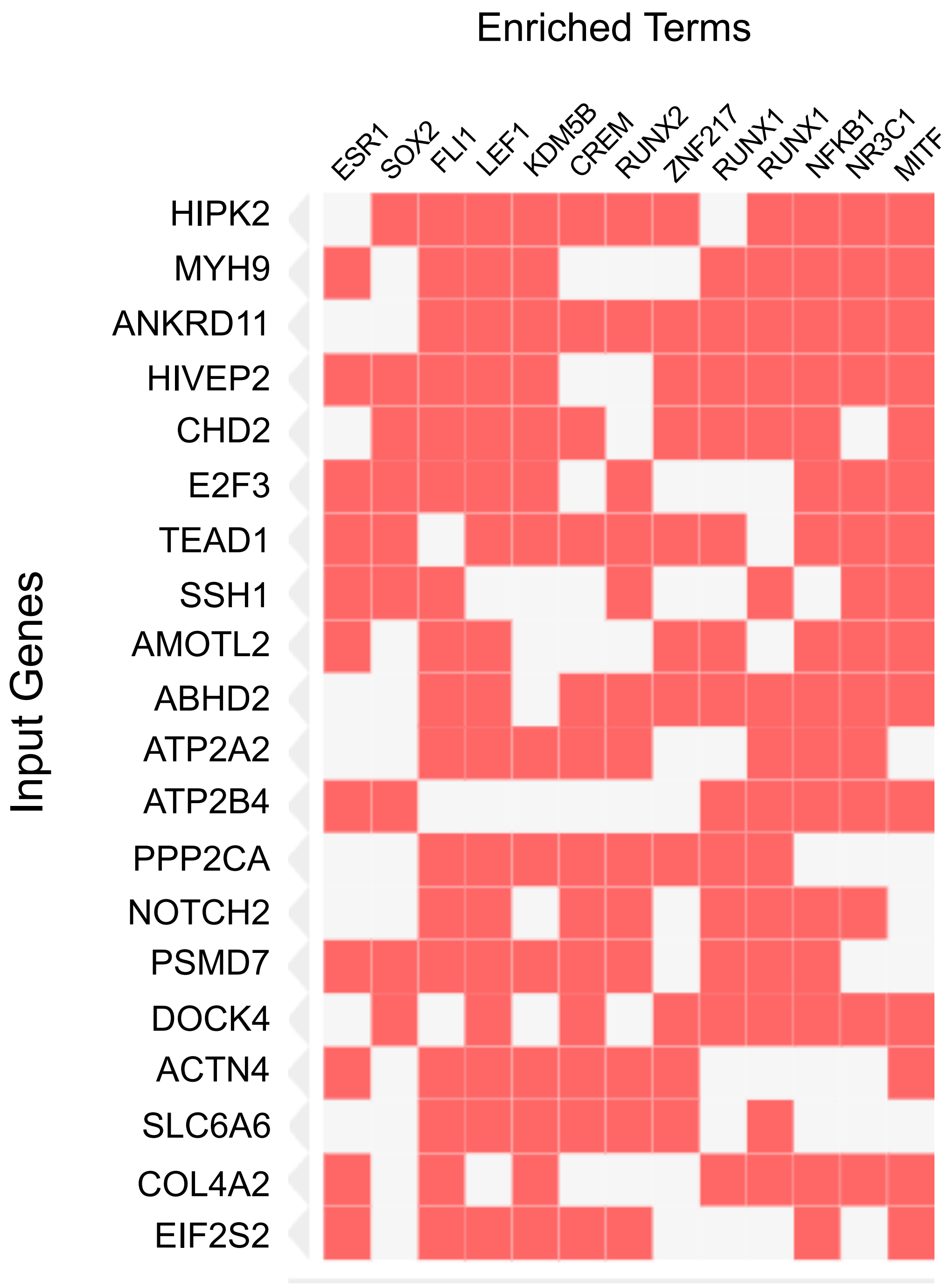


Supplemental Figure 5

A)

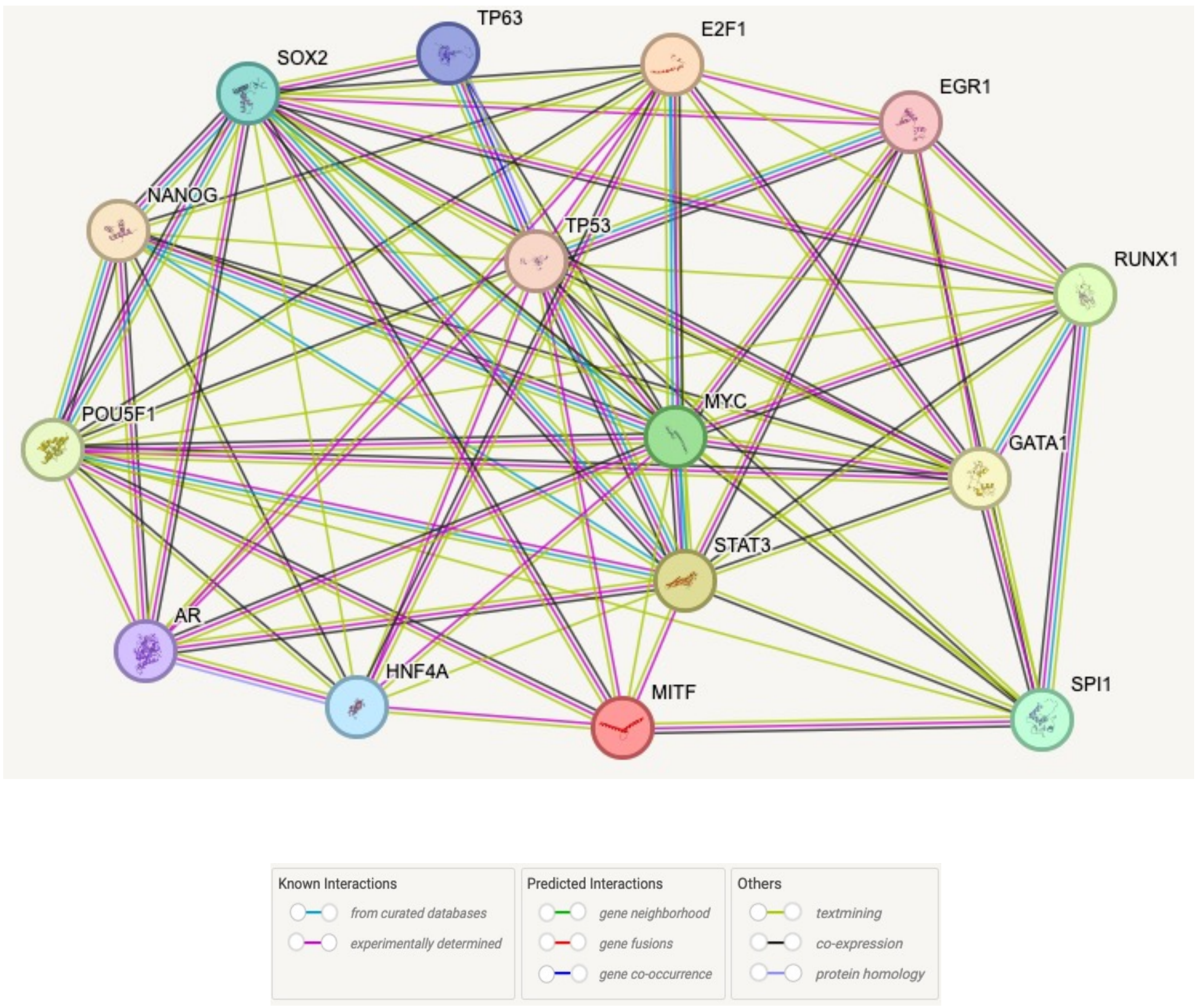


Supplemental Figure 6
